# Assessment of sample pooling for clinical SARS-CoV-2 testing

**DOI:** 10.1101/2020.05.26.118133

**Authors:** Sara B Griesemer, Greta Van Slyke, Kirsten St. George

## Abstract

Accommodating large increases in sample workloads has presented one of the biggest challenges to clinical laboratories during the COVID-19 pandemic. Despite the implementation of new automated detection systems, and previous efficiencies such as barcoding, electronic data transfer and extensive robotics, throughput capacities have struggled to meet the demand. Sample pooling has been suggested as an additional strategy to further address this need. The greatest concern with this approach in a clinical setting is the potential for reduced sensitivity, particularly the risk of false negative results when weak positive samples are pooled. To investigate this possibility, detection rates in pooled samples were evaluated, with extensive assessment of pools containing weak positive specimens. Additionally, the frequency of occurrence of weak positive samples across ten weeks of the pandemic were reviewed. Weak positive specimens were detected in all five-sample pools but failed to be detected in four of the 24 nine-sample pools tested. Weak positive samples comprised an average 16.5% of the positive specimens tested during the pandemic thus far, slightly increasing in frequency during later weeks. Other aspects of the testing process should be considered, however, such as accessioning and reporting, which are not streamlined and may be complicated by pooling procedures. Therefore, the impact on the entire laboratory process needs to be carefully assessed prior to implementing such a strategy.

## Introduction

The COVID-19 pandemic has presented numerous challenges to the health care industry in general and laboratory testing specifically. Not least among the latter has been a dramatic increase in sample testing load (1). Efforts to meet the demand have included increased use of automated instrumentation, multiplexing of molecular detection assays, streamlined testing protocols, as well as increasingly varied acceptable sample types, collection devices and transport media (2–4). Accommodating the testing workload and reagent shortage during the symptomatic pandemic wave was a significant undertaking (1). However, the more recent task, to test returning healthcare workers as well as patients returning for services such as nonessential surgery and other clinical procedures, has generated an even greater challenge. One proposed solution has been sample pooling (5–8), a method previously used for numerous other situations where large scale testing was needed (9–12).

When used as an epidemiological surveillance tool, acceptable parameters and limits may be quite different to those when the same system is applied in a clinical testing setting (6, 13–16). In the latter, the issue of detection sensitivity for every individual specimen becomes critical. Methods for pooling vary widely and testing large numbers of pooled samples is not possible without an inherent loss of sensitivity. However, there is the potential for more limited pooling, without loss of sensitivity. We sought to investigate to what extent this was possible, while maintaining the detection of weak positive samples.

## Methods

The CDC 2019 nCoV Real-Time RT-PCR Diagnostic Panel (17–19) was used throughout, with extraction on the bioMerieux EMAG^®^ (bioMerieux Inc, Durham, NC). For individual specimens, 110μL of Viral Transport Media (VTM, Regeneron, Rensselaer) or Molecular Transport Media (MTM, PrimeStore, LongHorn, San Antonio, TX) from upper respiratory swab, was added to 2mL NucliSens Lysis Buffer (bioMerieux) and extracted into 110μL of eluate.

The bioMerieux EMAG extraction system will accommodate a maximum volume of 3ml. Therefore, a maximum of nine samples (110 μL per sample) could be added to a 2ml lysis buffer tube without exceeding the maximum volume, while maintaining the same input volume per specimen. If the same extraction efficiency is maintained when nine samples are loaded in the tube as when one is loaded, and the eluate is still 110μL, theoretically the same total nucleic acid from each sample should be extracted. Provided there is no increase in the PCR inhibition of the pooled eluate, the detection sensitivity should therefore also still be the same.

Pool sizes of five and nine samples were tested, with each pool containing a single positive specimen. For specimen pooling, 110μL each of one positive, and either four (five-sample pool) or eight (nine-sample pool) negative specimens were added, at random, to 2mL of lysis buffer, in triplicate, extracted and eluted into 110μL of elution buffer.

In an initial experiment, strong and moderate positive samples in VTM were tested in each size pool. In a second experiment, four weak positive specimens in VTM and four weak positive specimens in MTM, were tested in both five and nine sample pools, in triplicate.

Additionally, the percentage of positive samples at different viral loads, as assessed by Ct value, was reviewed across nine weeks of the pandemic, to determine if these weak positive specimens comprise a significant component of total tested specimens and whether the proportion has changed over the course of the pandemic.

## Results

Pooling of samples with lower Ct values did not cause a loss of detection of any of the individual positive samples in either the five- or nine-sample pools (Table 1), although in one of the nine-sample pools, the detection became inconclusive rather than positive (specimen D). The Ct values of positive samples were increased by 0.4 to 1.51 when pooled with four negative samples. In contrast, pooling with 8 negative samples caused Ct increases ranging from 0.94 to 8.49.

**Table 1:** Effect of pooling SARS-CoV-2 positive NPS specimens with multiple negative NPS specimens.

| Pool Size | Positive Specimens<sup>#</sup> | N1* |  |  | N2* |  |  |
| --- | --- | --- | --- | --- | --- | --- | --- |
| | | Sample Ct | Pool Ct | $\Delta$ Ct | Sample Ct | Pool Ct | $\Delta$ Ct |
| 5 sample | A | 18.60 | 19.00 | 0.40 | 19.17 | 19.59 | 0.42 |
|  | B | 24.27 | 25.18 | 0.90 | 24.40 | 25.41 | 1.01 |
|  | C | 29.61 | 30.44 | 0.83 | 29.85 | 30.46 | 0.61 |
|  | D | 33.09 | 32.59 | -0.50 | 32.25 | 33.76 | 1.51 |
| 9 sample | A | 18.60 | 22.71 | 4.11 | 19.17 | 22.79 | 3.62 |
|  | B | 24.27 | 25.33 | 1.06 | 24.40 | 25.34 | 0.94 |
|  | C | 29.61 | 30.57 | 0.95 | 29.85 | 30.85 | 0.99 |
|  | D | 33.09 | 36.20 | 3.11 | 32.25 | 40.74 | 8.49 |

\* Virus real-time RT-PCR assay target
<sup>#</sup> in VTM

When weak positive samples were pooled with four or eight negative samples (Table 2), the positive samples were still all detected in five-sample pools, whether they were in VTM or MTM transport media. When combined in nine-sample pools, detection was more adversely affected for samples in VTM than those in MTM. For samples in MTM, one of three replicates for one sample, failed to be detected in a pool of nine samples. In contrast, for weak positive specimens in VTM transport media, nine-sample pools caused multiple replicates to return negative results.

**Table 2:**
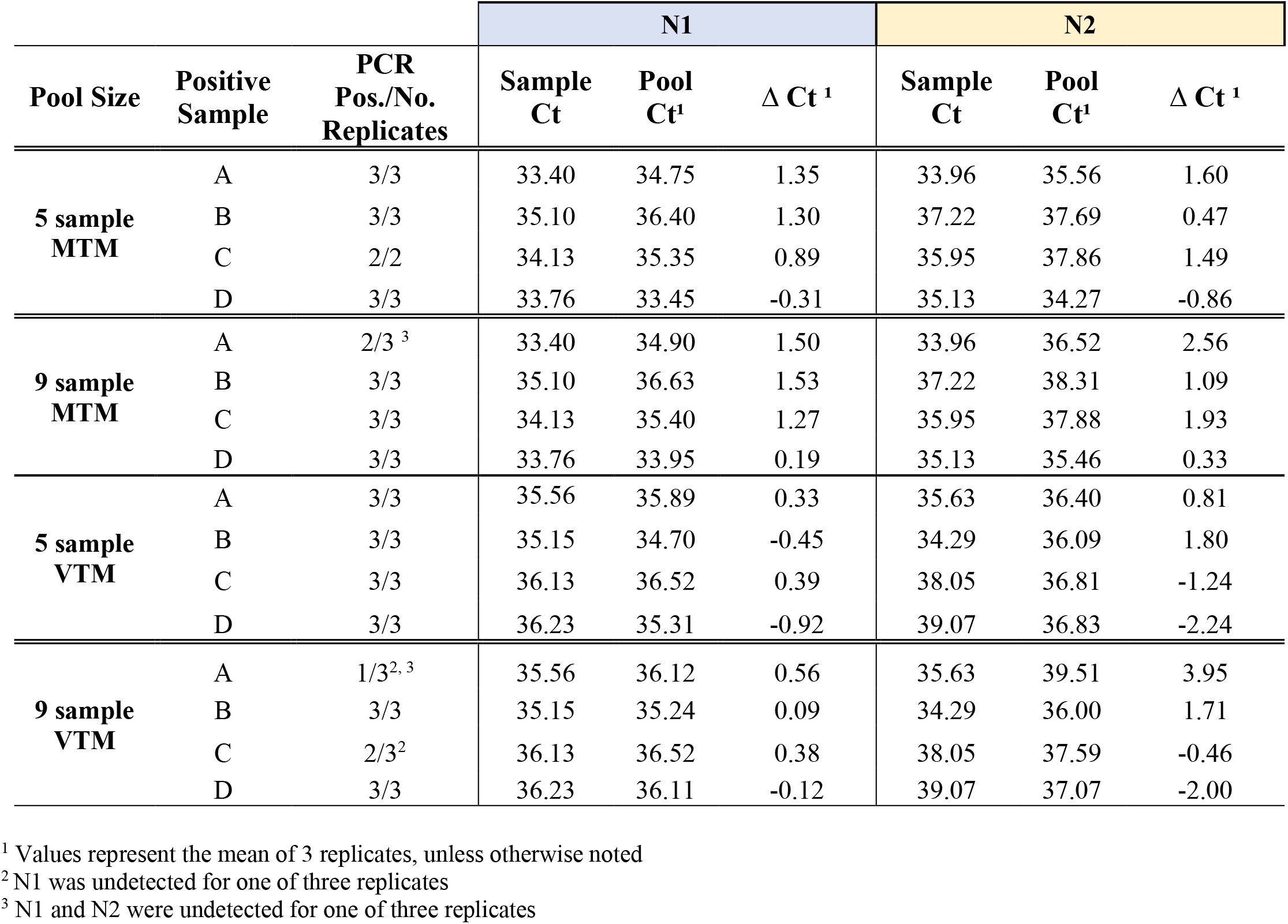
Effect of pooling low-titer SARS-CoV-2 positive NPS specimens with multiple negative NPS specimens.

We then sought to assess what component of the total specimens tested are comprised of these weak positive specimens, to evaluate how much of an impact pooling might have overall on testing sensitivity across positive patient detection in the pandemic. Further, to assess the positivity rate during the months since the onset of the pandemic in New York, since pooling strategies are not efficient unless sample positivity is low. Despite the large range in number of specimens tested per day from late February through mid-May (Figure 1), the percentages of specimens with viral loads ranging from very strong (Ct < 20), strong (Ct 21-25), moderate (Ct 26-30), weak (Ct 31-35) and very weak (Ct 36-40) remained remarkably constant, with the exception of those in the very weak range which increased slightly during the last five weeks. Overall, this weak positive sample type constituted an average 16.5% of positive specimens, a substantial proportion of the positive specimens received for testing.

**Figure 1.**
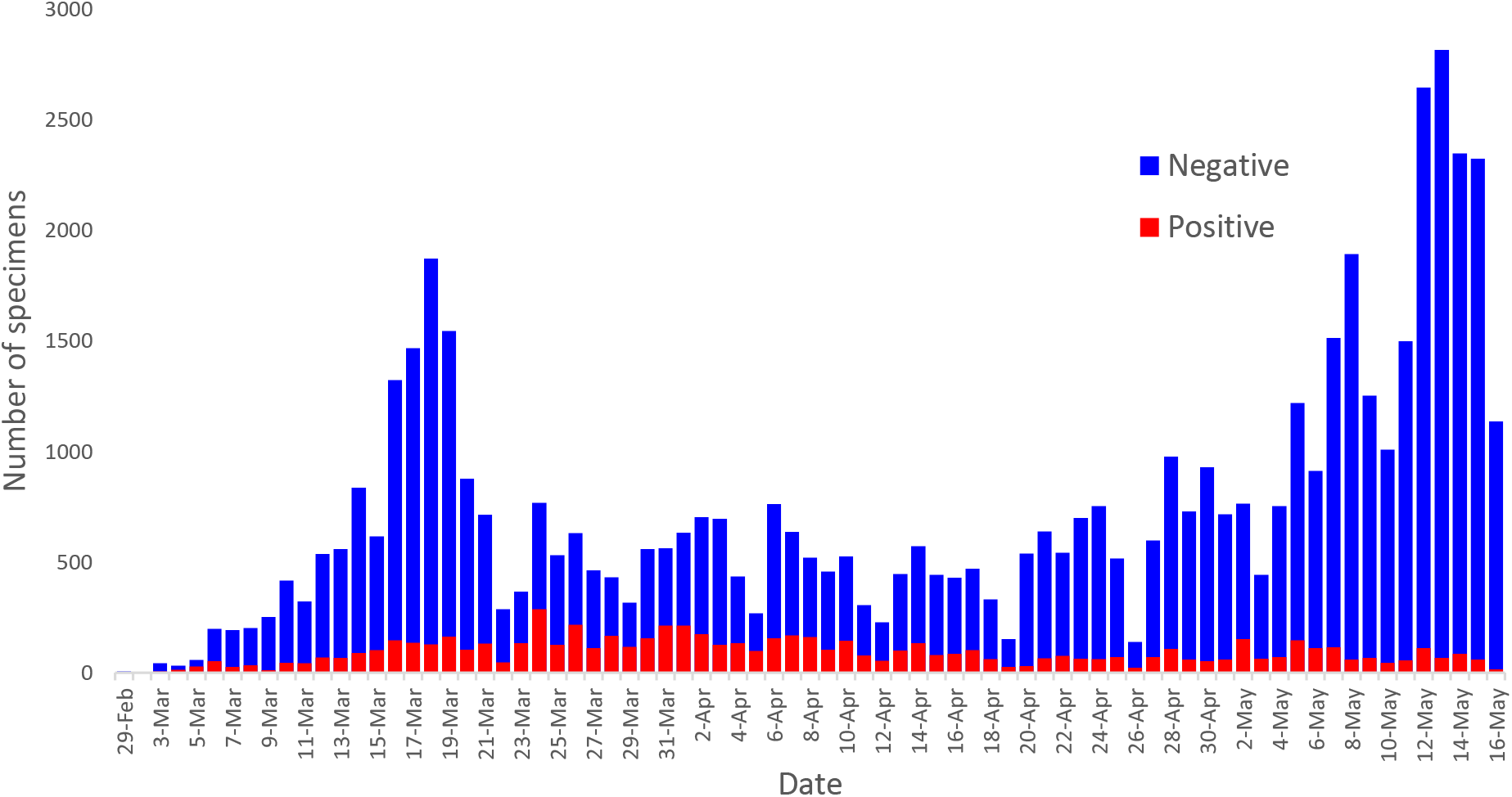
Number of specimens tested per day at the Wadsworth Center for SARS-CoV-2, from February 29 to May 16, 2020. Negative specimens in blue bars, positive specimens in red bars.

Positivity rates among samples received at this facility rose to almost 30% during March and has continually dropped since early April, remaining below 5% since 10^th^ May and below 1.5% since May 16.

## Discussion

As the pandemic evolves, despite case counts, hospitalization rates and fatalities decreasing in some areas, laboratories continue to face new challenges. Workloads have increased with, for example, requirements to perform repeated surveillance screening of asymptomatic health care workers and testing of patients undergoing elective procedures where there is a risk of aerosol production. These policies have pushed test numbers beyond those encountered even at the height of the pandemic wave. Suggestions to help manage the load have included pooling of samples to enhance throughput capacity. When being considered for application in a clinical testing environment, of greatest concerns is the potential for this strategy to increase the false negative rate. With that, the greatest risk of false negative results is with weak positive samples. To investigate the potential limits of pooling in this situation, this study focused on pooling weak positive samples in relatively small size sample pools. Had these been successful, larger pools would have been attempted.

There are multiple methods for pooling samples, some of which carry an inherent risk of reducing test sensitivity and some of which do not. The method described in this paper, where the same volume of each sample is added to lysis buffer as would normally be added if the sample were being tested individually, does not theoretically adversely affect sensitivity, as long as extraction efficiency is maintained and the level of PCR inhibition is not increased by the additional load through the extraction device. When the data was analyzed, despite some minor shifts in Ct values, no loss of detection was observed in any of the five-pool experiments, for samples that had been collected in either VTM or MTM. However, when the same weak positive samples were pooled with 8 negative samples rather than 4, to create pools of 9, detection failures were observed. In three of the 9-sample VTM pools and one of the 9-sample MTM pools, this larger pool size resulted in a complete loss of detection. Whether the less frequent occurrence in MTM samples compared to VTM samples is significant is difficult to say based on this limited data.

The use of pooling as a throughput enhancement strategy is only efficient if the positivity rate in the samples being tested is low enough that a minimal number of pools will test positive, otherwise, multiple pools will have to be deconvoluted for retesting of individual samples. The optimal or maximum positivity level at which pooling starts to become efficient, depends on the pool size being used. For pool sizes as small as 5 samples, this maximum positivity level is considerably higher than that for very large pool sizes that are sometimes suggested for large scale epidemiological screening studies. For example, at a positivity rate of 1% and a pool size of 5, on average, only 1 in every 20 pools will be positive and need to be deconvoluted for retesting. Therefore, for every 100 samples, testing could be achieved with a total of 25 tests (20 pools and one deconvoluted pool). We noted that the positivity rate in our own lab is now approaching 1% and therefore such a strategy may be efficient for extraction and detection. Moreover, as the pandemic has progressed, there has been an increasing proportion of samples in the weak positive range, and therefore major consideration must be given to the issue of detection sensitivity for these weak samples. It must be noted however, that extraction and detection are not the only components of the laboratory operation and a pooling strategy does not enhance other aspects and may in fact create complications.

Processes for specimen receiving and accessioning, as well as those for result data management and reporting, are not reduced by pooling strategies. These procedures may be complicated by sample pooling, especially for electronic data transfer programs and laboratory information systems. A large increase in test load facilitated by a pooling strategy, may create serious workload bottlenecks for these and other areas of the operation. Therefore, the global implications of a pooling strategy need to be carefully assessed, especially in the clinical testing environment, before implementation.

**Figure 2.**
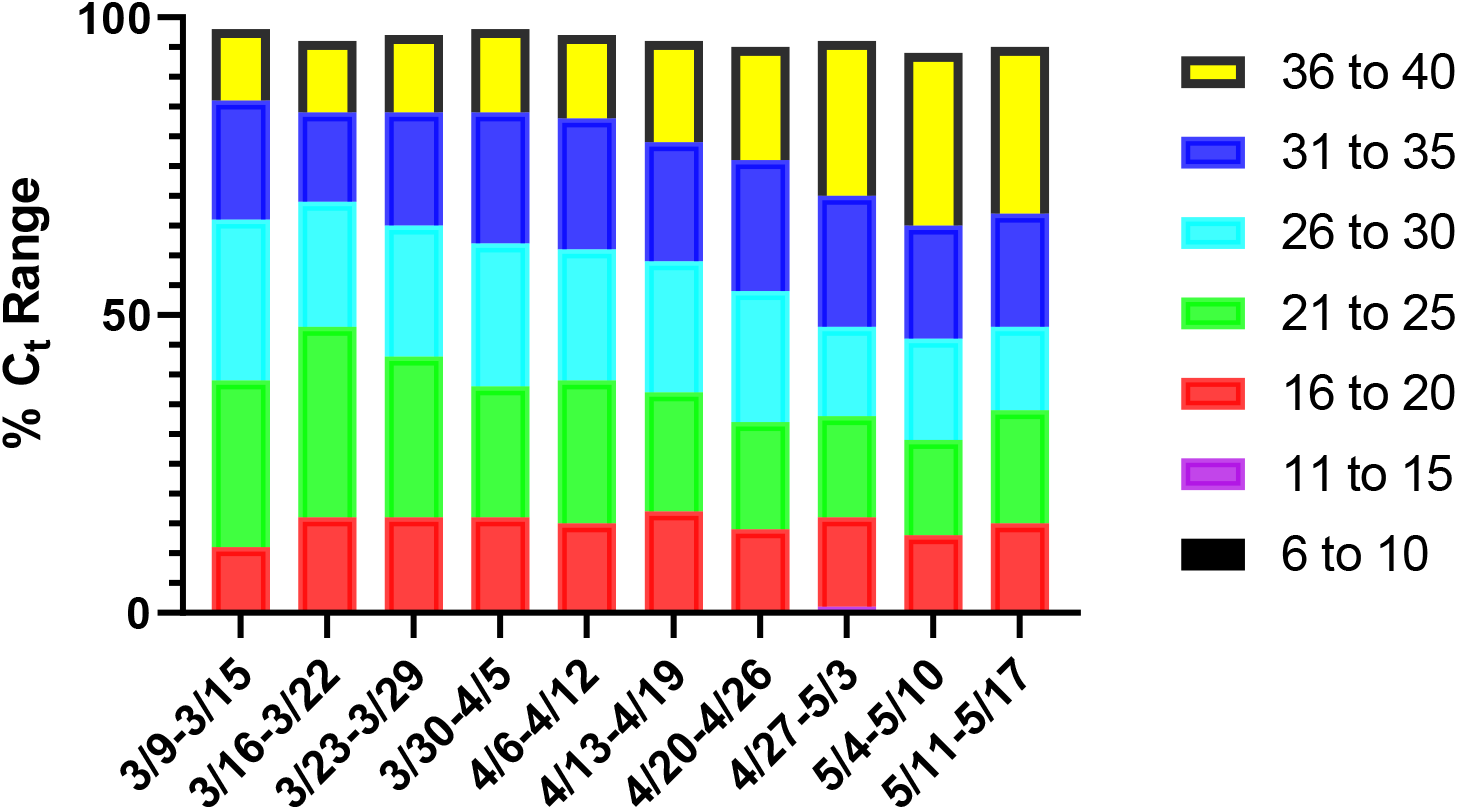
Percentage of specimens in each category of viral load, as approximated by Ct range, by week, from week starting 3//9/20 to week starting 5/11/20. Ct values are the average of the N1 and N2 value for each specimen.

## Acknowledgements

The authors thank Kamran Zamani for assistance with data management and the provision of Figure 1. They also thank Jennifer Laplante, Rene Hull and Steven Zink for specimen selection and retrieval. We thank the Wadsworth COVID response team for months of dedicated, skilled testing and careful specimen archiving that provided the well-characterized samples accessed for this work.

